# The effects of social environment and the metapleural gland on disease resistance in acorn ants

**DOI:** 10.1101/2021.12.14.472662

**Authors:** Joseph T. Scavetta, Sarah F. Senula, Daniel R. Crowell, Farzana Siddique, Jennifer F. Segrest, Olseun A. Dairo, Lindsey U. Nguyen, Mathew S. Pekora, Svjetlana Vojvodic Kruse

## Abstract

Eusocial species differ in living conditions when compared to solitary species primarily due to the dense living conditions of genetically related individuals. Consequently, these crowded conditions can induce a high rate of pathogen transmission and pathogen susceptibility. To resist an epidemic, individuals rely on sets of behaviors, known as social immunity, to decrease pathogen transmission among nestmates. Alongside social immunity, ants have a pair of secretory metapleural glands (MG), thought to secrete antimicrobial compounds important for antisepsis, that are believed to be transferred among nestmates by social immune behaviors such as grooming. To investigate the effects of social immunity on pathogen resistance, we performed a series of experiments by inoculating acorn ants *Temnothorax curvispinosus* with a lethal spore concentration of the entomopathogenic fungus *Metarhizium brunneum*. After inoculation ant survival was monitored in two environments: solitary and in groups. Additionally, the MG role in pathogen resistance was evaluated for both solitary and grouped living ants, by sealing the MG prior to inoculations. Individuals within a group environment had a higher survival compared to those in a solitary environment, and individuals with sealed glands had significantly decreased survival than ants with non-sealed-MG in both solitary and social environments. We observed the lowest survival for solitary-sealed-MG individuals. Although sealing the MG reduced survival probability, sealing the MG did not remove the benefits of grouped living. We show here that social living plays a crucial role in pathogen resistance and that the MG has an important role in pathogen resistance of individual *T. curvispinosus* ants. Although important for an individual’s pathogen resistance, our data show that the MG does not play a strong role in social immunity as previously believed. Overall, this study provides insights into mechanisms of social immunity and the role of MG in disease resistance.

## Introduction

There are many benefits of social living, such as shared resource and protection from predators, but the inherent colony structure could also be thought to increase pathogen spread among individuals [1]. Historically, the infection rate across a group of individuals is assumed to be linearly dependent on the density of hosts and pathogens [2]. Pathogen transmission is increased with contact, and this would certainly be true for dense colonies, especially observed in the insect order Hymenoptera, where individuals are constantly in contact by exchanging food and information [3,4]. Furthermore, a high interaction rate among genetically related individuals has the potential to increase both the spread and virulence of the pathogen [1,5]. However, eusocial insects such as ants, bees, termites and wasps have an ecologically high success rate by inhabiting many types of ecosystems, and often dominate their surrounding environments [6,7].

The large prominence of social insects must indicate evolutionary adaptations to the increased pathogen risk found with eusocial living [8]. Importance of individual immunity for disease resistance is well documented [9,10], but the growing body of literature demonstrates the importance of group immune responses primarily observed in social insects [5,8,11,12,13,14,15]. Many adaptations have been suggested, encompassing behavioral, spatial, and physiological solutions that often work on a colony level, generally known as *social immunity* [16]. This encompasses several different behaviors such as grooming, corpse removal of infected cadavers [17,18], removal of infectious waste from hives [19], and task partitioning as individuals age, known as temporal polyethism [20]. Furthermore, some social insects use antimicrobial compounds either directly against the pathogen, such as in oral secretions from termites [21], or within their nests, such as plant sap in a form of propolis in honey bees [11,22]. Along with pathogen transmission prevention, social insects can provide nestmates with contact immunization, in which a non-infected individual receives the pathogen to increase resilience in potential future pathogen contact [23,24,25]. Similarly, pharaoh ants prefer a pathogen-exposed nest over noninfected nests, increasing the immune priming of the whole colony [26]. The presence of social immunity is thought to have arisen to maximize inclusive fitness of each individual, opening many altruistic traits and actions that are vital for the colony as a whole and are comparable to an organism level immune system [27].

Along with these adaptations, ant species have a pair of secretory metapleural glands (MG) found in the posterior of the mesosoma; these glands likely have evolved once within the Formicidae family [28]. Among the proposed functions of the gland, secretions of pheromones [29], defensive chemicals [30], and antibiotics [31,32,33,34] have been observed from varying ant species, but the primary function of the MG is believed to be antisepsis [28]. The evolution of a gland specialized in hygienic secretions may be an important aspect of pathogen resistance among highly social individuals and may increase resistance among nestmates [28]. However, loss of the MG has occurred within multiple species, notably within the Formicinae subfamily [35], raising questions about the gland’s true function. Though increased pathogen susceptibility is predicted, this was not observed in *Polyrhachis dives*, indicating other mechanisms of pathogen defense such as increased self-grooming [36].

In this study, we examined the high-level effects of social immunity on the survival probability of *Temnothorax curvispinosus* infected with a generalist fungal pathogen, *Metarhizium brunneum*. We analyzed the survival probabilities of *T. curvispinosus* when in groups compared to when in isolation. Following this, we investigated the MG’s function in terms of the antisepsis hypothesis, specifically if the gland was important for pathogen resistance at the individual level as well as the group level. To explore this, we developed and tested an approach to sealing the MG that was used to compare the survival probability of infected ants without functioning MG to the ants that have fully functioning glands. Finally, we compared the MG’s role in ant survival between two environments: solitary and group living.

## Material and methods

### Host maintenance

Acorn ant, *Temnothorax curvispinosus*, colonies were collected in Spring 2015 from Scotland Run Park, Clayton, New Jersey (39.656 -75.052) and maintained at Rowan University. Each colony was stored in a Fluon-coated (BioQuip Products, Rancho Dominguez, CA, USA) 15 cm diameter Petri dish with a glass water tube sealed with a cotton wool. Colonies were fed weekly with half a cricket and one drop of diluted honey that was placed on approximately 1 cm square paper towel to prevent drowning. Large single queen colonies were used only once in each experiment.

### Pathogen maintenance and inoculum preparation

The fungus *Metarhizium brunneum* (strain ARSEF 1095), previously named *Metarhizium anisopliae* [37], was originally isolated from *Carpocapsa pomonella* (Lepidoptera: Olethreutidae) in Austria and was ordered from the USDA-ARS Collection of Entomopathogenic Fungi Cultures in Ithaca, New York, USA. This strain was maintained on Sabouraud Dextrose agar (SDA) plates and were transferred to new SDA plates monthly.

A new conidiospore inoculum was prepared on the day of ant’s inoculation experiment by covering the Petri dish plate with a solution of 0.05% Triton-X (prepared from a Triton X-100 (Sigma-Aldrich, St. Louis, USA)). After the fungal conidiospores were adequately mixed in Triton-X solution, a 1-mL micropipette was used to extract and transfer the solution into a 15 mL centrifuge tube. In order to eliminate other fungal particles, the solution was allowed to settle for approximately 30 min after which time the upper layer of the solution that contained only conidiospores was transferred into a new 15 mL tube. The conidiospore concentration of the acquired suspension was then determined with a hemocytometer (Neubauer-improved counting chamber) and diluted to a final conidiospore concentration needed for each of the experiments outlined below.

### A. Test of dosage response and group living on disease resistance

#### I. Dosage response

For this experiment we used six single queen colonies with over 100 worker ants. The colonies were cooled at 4°C to slow the ant movement and kept for 5 min on ice packs while 100 ants were transferred from a source colony into a temporary Petri dish. From there ants were evenly divided into three inoculation treatments: i) conidiospore inoculum 1 × 10^5^ conidiospore/mL; ii) conidiospore inoculum 1 × 10^7^ spore/mL; and iii) sham inoculated with 0.05% Triton-X solution totaling 32 ants per treatment per colony. For each ant in each treatment, the ant was submerged within the corresponding solution for 5 seconds and set upon a paper towel to dry. Immediately after drying, the ant was placed into a 1.5 mL microcentrifuge tube that was then capped with moist cotton wool. Ants remained in individual microcentrifuge tubes for 24 hours to ensure infection and to control for allogrooming of conidiospores that would result in variable ant infections. Following the 24-hour isolation period, ants from each colony and inoculation treatment were moved for the last time: with half of the ants (per colony/treatment) being moved into a glass tube in a group of 16 ants, and the other half (16 ants) were kept by in individual glass tubes. All tubes, with group and individual ants, were sealed with moist cotton wool plugs that also provided water to the ants. Small paper towel cuttings (0.5 × 0.5 cm) dipped in diluted honey were added 24 hours after the ants were placed into the tubes. Survival was checked and recorded daily for a total duration of 12 days post-inoculation.

#### II. Survival in groups vs. solitary environment

Six single queen colonies containing over 110 workers were cooled at 4°C until movement slowed; colonies were then placed on an ice pack while 110 workers were randomly selected from each colony into colony specific Fluon-coated petri dishes with water and diluted honey.

Ants from each colony were divided into two inoculation treatments: 1. *M. brunneum* conidiospore inoculated (1 × 10^8^ conidiospores/mL) (n=48 per colony), and 2. sham inoculated (0.05% Triton-X solution) (n=62 per colony). For each ant in each treatment, the ant was submerged within the corresponding solution for 5 seconds and set upon a paper towel to dry. Immediately after drying, the ant was placed into a microcentrifuge tube that was then capped with moist cotton. Ants remained in individual microcentrifuge tubes for 24 hours.

After the 24-hour isolation period, ants from each inoculation treatment were divided into social treatments: 1. individual glass tubes (n = 26 per inoculation treatment per colony), 2. group glass tubes of 13 ants (n = 26 sham per colony, 13 inoculated per colony), and 3. group glass tubes containing 5 conidiospore inoculated and 8 sham inoculated ants (n = 13 per colony). Moist cotton was used to seal the tubes and provide water to the ants. Small paper towel cuttings dipped in diluted honey were added 24 hours after the ants were placed into the tubes. Survival was checked and recorded daily for a total duration of 21 days post-inoculation.

### B. Metapleural gland impact on disease resistance

#### I. Effects of solitary environment, CO_2_, and superglue on ant survival

Three single queen colonies containing over 120 workers were sedated using carbon dioxide until movement slowed; 120 workers were randomly selected from each colony into colony specific Fluon-coated Petri dishes. Ants from each colony were divided into four treatments: 1. No treatment (n ≈ 30 per colony), 2. exposure to CO_2_ for (4 min) (n ≈ 30 per colony), 3. exposure to superglue onto the thorax (n ≈ 30 per colony), and 4. sealing of the metapleural glands (MG) with superglue while sedated with CO_2_ (n ≈ 30 per colony).

Following the treatments above ants from each colony and treatment were then split into two inoculation treatments: 1. pathogen *M. brunneum* conidiospore inoculated (1 × 10^8^ conidiospores/mL) (n ≈ 15 ants per treatment per colony), and 2. sham inoculated (0.05% Triton-X solution) (n ≈ 15 ants per treatment per colony). For each ant in each treatment, the ant was submerged within the corresponding solution for 5 seconds and set upon a paper towel to dry. Immediately after drying, the ant was placed into a microcentrifuge tube that was then capped with moist cotton. Ants remained in individual microcentrifuge tubes while survival was checked and recorded daily for a total duration of 12 days post-inoculation.

#### II. Solitary vs. group environment with open and closed metapleural gland

Three single queen colonies containing over 96 workers were sedated using carbon dioxide until movement slowed; 96 workers were randomly selected from each colony into colony specific Fluon-coated petri dishes. Ants from each colony were divided into two sealing treatments: 1. non-sealed metapleural gland, and 2. sealed metapleural gland. Ants that had their MG sealed were sedated with CO_2_ while superglue was placed over their metapleural glands.

Ants from each sealing treatment were then divided into two inoculation treatments: 1. *M. brunneum* conidiospore inoculated (2 × 10^8^ conidiospores/mL), and 2. sham inoculated (0.05% Triton-X solution). For each ant in each treatment, the ant was submerged within the corresponding solution (conidiospore presence/absence) for 5 seconds and set upon a paper towel to dry. Immediately after drying, the ant was placed into a microcentrifuge tube that was then capped with moist cotton. Ants remained in individual microcentrifuge tubes for 24 hours to reduce conidiospore removal by allogrooming and to ensure conidiospore germination. Following the 24-hour isolation period, ants from each colony were divided into two social treatments consisting of either solitary or group glass tubes. The solitary tubes were divided into four combinations: 1. conidiospore inoculated and gland-sealed ants, 2. conidiospore inoculated and non-sealed ants, 3. sham inoculated and gland-sealed ants, and 4. sham inoculated and non-sealed ants. The group tubes were also divided into four combinations: 1. all conidiospore inoculated and gland-sealed ants, 2. all sham inoculated and gland-sealed ants, 3. four conidiospore inoculated and gland-sealed ants with eight sham inoculated and non-sealed ants, and 4. four conidiospore inoculated and non-sealed ants with eight sham inoculated and gland-sealed ants. All ants were moved to the respective tubes and moist cotton was used to seal the tubes and to provide water to the ants. Survival was checked and recorded daily for a total duration of 12 days post-inoculation.

A time replicate was conducted 79 days after the first; all methods described above were repeated with four new single queen colonies.

### Statistical Analysis

All statistical analyses were performed using RStudio v. 1.0.136 with R v. 3.3.2. All survival data were analyzed using the survival package [38], making use of the Kaplan–Meier estimator. Significance of survival data was determined by using the log-rank test on survival probability estimators. All survival probabilities are reported as the 95% log-log confidence intervals. Alpha was corrected from α = 0.05 for multiple comparisons using Bonferroni Correction on each experiment. For the dosage response, alpha was adjusted to 0.0167. For the investigation of group versus solitary conditions, alpha was adjusted to 0.0033. For the metapleural gland experiments, alpha was adjusted to 0.0018.

## Results

### Dosage response and immune priming

The *M. brunneum* conidiospore concentration treatment had an effect on the survival of exposed ants (See supplemental). The lowest pathogen concentration tested was 10^5^ conidiospores/mL that had no significant difference on ant survival compared to sham inoculated ants (χ2 = 1.05, d.f. = 1, p = 0.305). Ants inoculated with 10^7^ conidiospores/mL had a significant decrease in survival when compared to sham inoculated ants (χ2 = 7.80, d.f. = 1, p = 0.005), and to 10^5^ conidiospore/mL inoculated ants (χ2 = 12.69, d.f. = 1, p < 0.001). Host survival was recorded for twelve days and within that time the following survival probabilities were recorded for each treatment: sham inoculated ants survived within 0.93 to 1.00, ants exposed to 10^5^ conidiospores/mL survived within 0.97 to 1.00, and ants exposed to 10^7^ conidiospores/mL survival probability within 0.78 to 0.93.

### Effects of social grouping on survival

Social environment vs. host isolation (solitary) had a significant effect on the survival of ants that were inoculated with *M. brunneum* (**Fig 1**). Furthermore, high survival rate in a control treatment with a sham inoculated ants validates our experimental protocol for both condition with the solitary treatment not differing from the group treatment for either experiment 1 (*χ*^*2*^ = 3.07, *d*.*f*. = 1, *p* = 0.08) or experiment 2 (*χ*^*2*^ = 0.27, *d*.*f*. = 1, *p* = 0.605), that collectively had a probability of survival within 0.94 to 1.00 for a duration of twelve days. In both experiments (1 and 2), all sham inoculated treatments differed from the treatments involving pathogen inoculations, for solitary (*χ*^*2*^ = 255.1, *d*.*f*. = 1, *p* <0.001) and group treatments (*χ*^*2*^ = 64.71, *d*.*f*. = 1, *p* <0.001). As expected, inoculated ants living within a group had a significantly higher survival probability than inoculated ants in solitude (*χ*^*2*^ = 13.45, *d*.*f*. = 1, *p* < 0.001). Within twelve days, inoculated ants in a group treatment had a probability of survival within 0.35 to 0.57, whereas inoculated ants in solitude had a much lower survival probability within 0.12 to 0.24.

**Fig 1.**
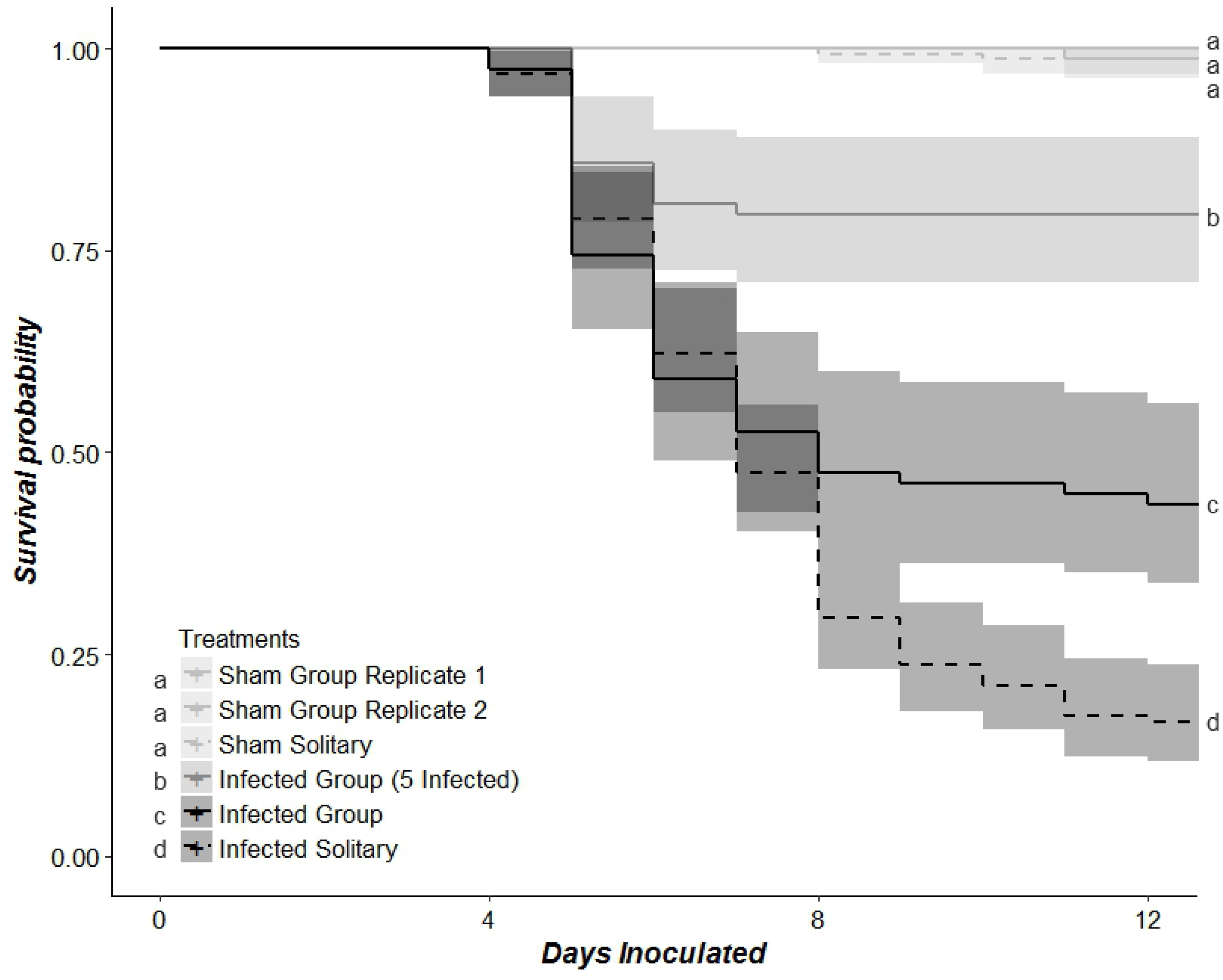
The effects of inoculation of *M. brunneum* conidiospores and the effects of social conditions on the survival of *T. curvispinosus*. Solid lines depict ants in group conditions, dashed lines depict ants in solitary conditions. Darker shading indicates spore inoculated ants. Sham inoculated treatments do not differ from one another regardless of social condition, though they do differ from all pathogen inoculation treatments. Spore inoculated ants differ depending on social conditions, where solitary ants have the lowest survival probability in a timeframe of twelve days. Different letters show significance at α = 0.003334.

### Effects of metapleural gland on survival

To validate our experimental protocol, ants were subjected to CO_2,_ a glue treatment on their back, or sham inoculation, and all ants were kept in individual tubes by themselves. These two treatments did not differ from ant’s survival that were only sham inoculated (CO_2_: *χ*^*2*^ = 0, *d*.*f*. = 1, *p* = 1.000; Glue: *χ*^*2*^ = 3.140, *d*.*f*. = 1, *p* = 0.076) (**Fig 2**). The survival probabilities at the end of twelve-day experiment, of sham inoculated ants with no further treatment with CO_2_ and glue was as follows: sham 1.00, CO_2_ 1.00, and glue 0.86 to 1.00. The survival probabilities of pathogen inoculated ants with no further treatment, CO_2_ treatment, and glue treatment was: pathogen only 0.18 to 0.46, CO_2_ 0.24 to 0.53, and glue 0.20 to 0.48.

**Fig 2.**
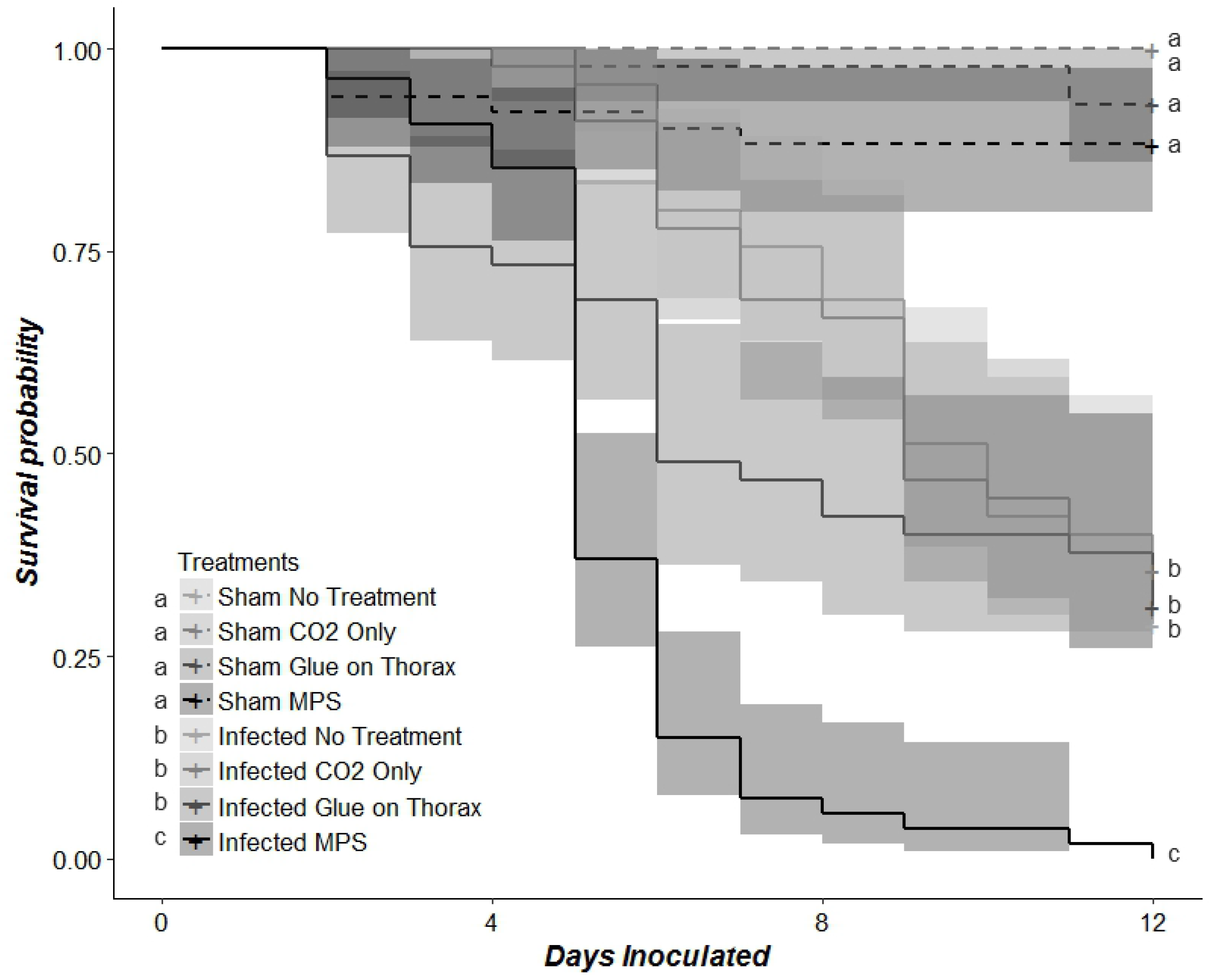
The effects of CO_2_, superglue, and the metapleural gland on the survival of *T. curvispinosus*. Solid lines depict ants inoculated with *M. brunneum* conidiospores, dashed lines depict ants sham inoculated. Sham inoculated treatments do not differ from one another regardless of CO_2_, glue, or the sealing of the metapleural glands. All sham and pathogen inoculated ants differ from one another in survival probability for the timeframe of twelve days. Spore inoculated ants do not differ from each other by CO_2_ or glue; however, the spore inoculated ants with their metapleural glands sealed differ significantly from all other treatments. Different letters show significance at α = 0.0018.

Sham inoculated ants with MG-sealed had a survival probability of 0.80 to 0.98 and did not significantly differ in survival compared to sham inoculated ants with no further treatments (*χ*^*2*^ = 5.58, *d*.*f*. = 1, *p* = 0.018). However, ants that had their MG-sealed and were pathogen inoculated had a survival probability of 0 in the timeframe of twelve days and this was significantly different when compared to pathogen inoculated ants with no further treatments (*χ*^*2*^ = 56.46, *d*.*f*. = 1, *p* < 0.001,) showing that MG is indeed important during pathogen exposure **(Fig 2)**.

Sealing the MG had a significant effect on the survival of solitary ants that were inoculated with *M. brunneum* conidiospores, but not for sham inoculated ants **(Fig 3A)**, and the same trend was observed for ants kept in groups **(Fig 3B)**. Conidiospore inoculated ants in solitary conditions differed based on the sealing of the MG (*χ*^*2*^ = 75.96, *d*.*f*. = 1, *p* <0.001), with sealed-solitary ants having the lowest survival in the experiment with a probability of 0 survival within twelve days. Non-sealed MG solitary infected ants had a slightly higher survival probability of 0.01 to 0.09.

**Fig 3.**
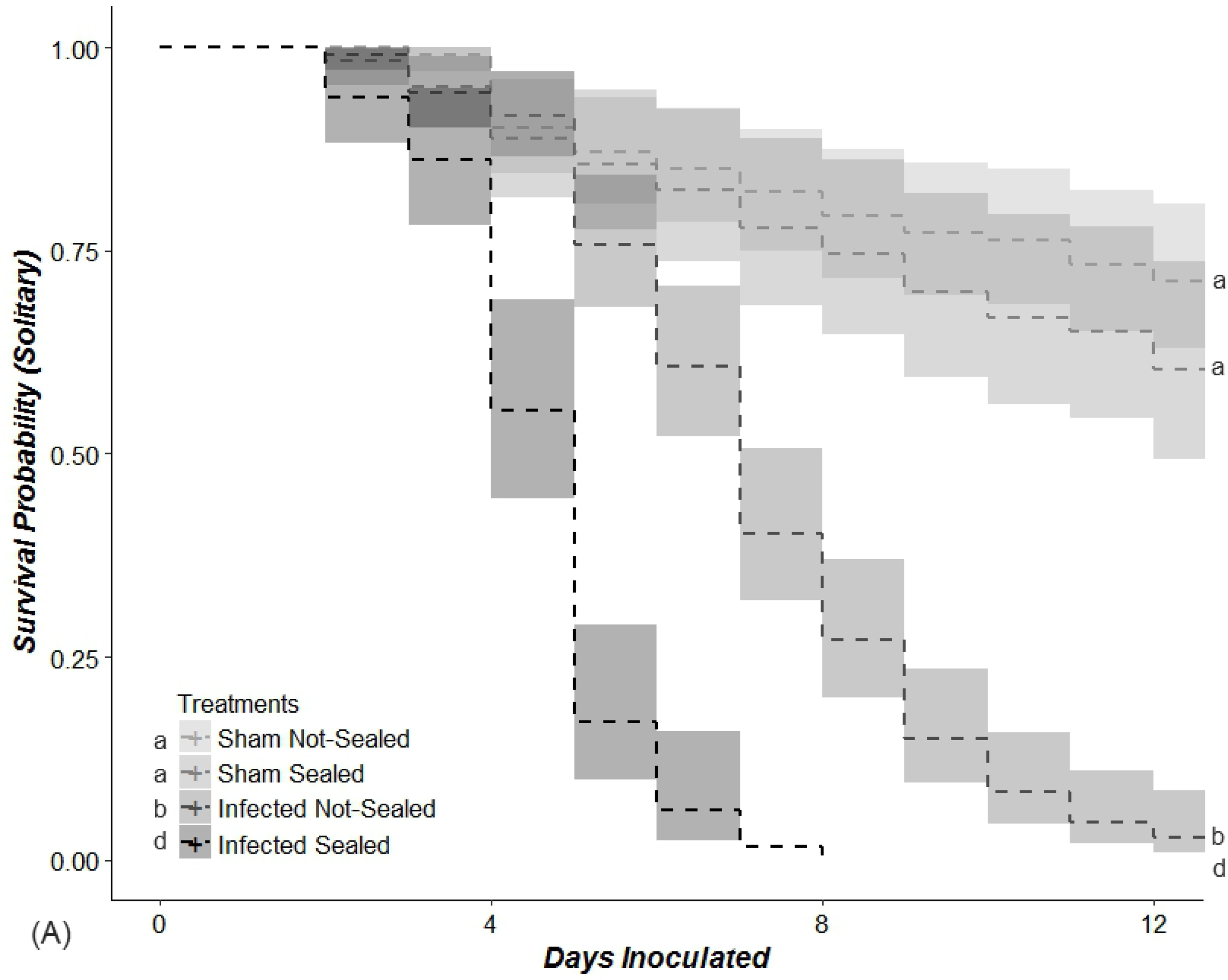
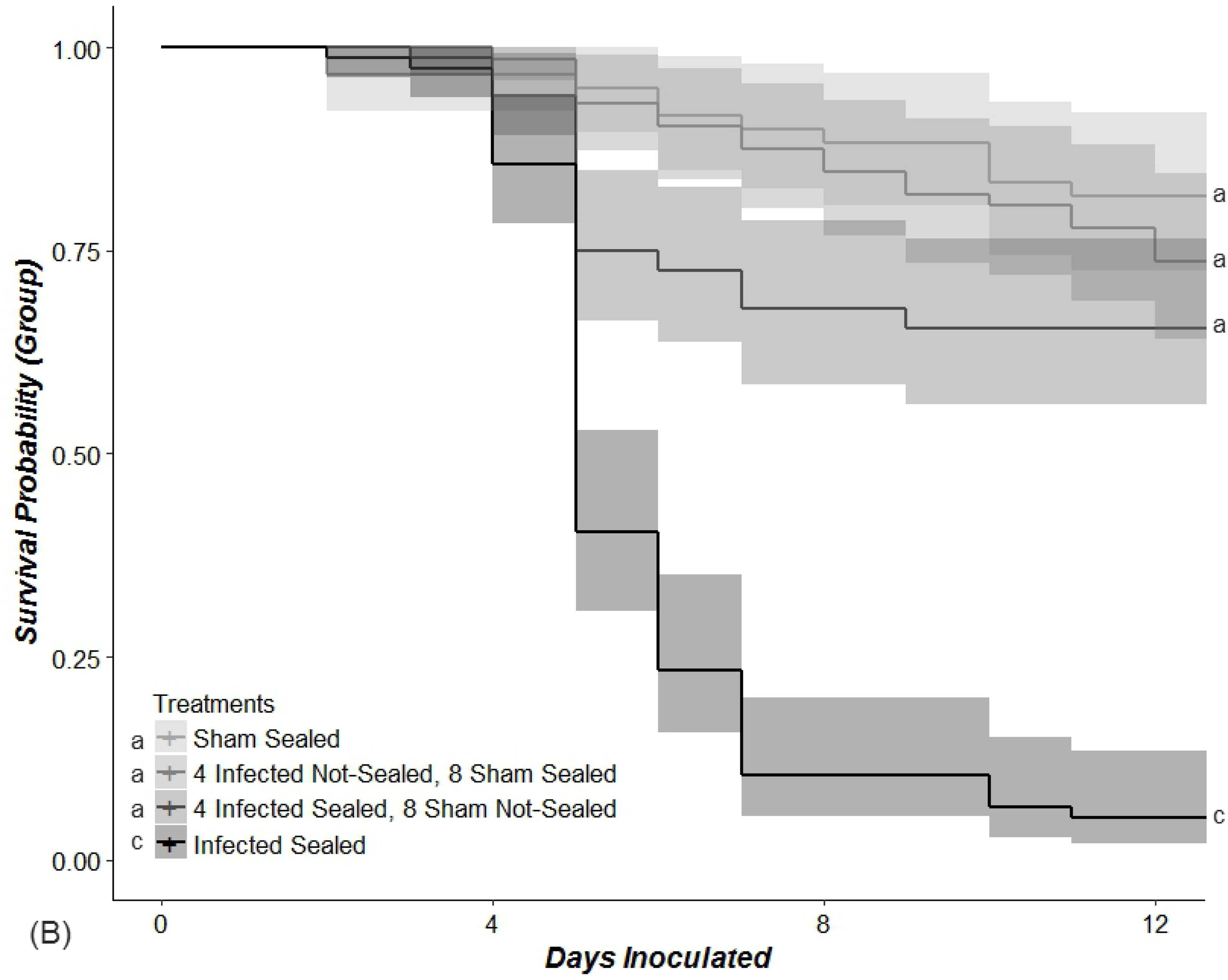
The effects of the metapleural gland with group conditions on the survival of *T. curvispinosus*. (A) Shows ants in solitary conditions, while (B) shows ants in group conditions. Sham inoculated ants did not differ from one another regardless of the sealing of the metapleural gland or their social conditions. Groups in which sham and pathogen exposed ants with differing sealing treatments were mixed did not differ from the sham inoculated treatments. Spore inoculated ants with and without a sealed metapleural gland differed from all sham treatments, but also differed significantly from each other as sealed ants had a lower survival probability. Spore inoculated ants with a sealed metapleural gland also differed depending on social conditions, where those in a group had a higher survival than those in solitary. Different letters show significance at α = 0.0018.

Survival of infected ants in two tested environments had the most pronounced survival differences with the MG-sealed infected ants in solitary conditions differing significantly from MG-sealed conidiospore inoculated ants in group conditions (*χ*^*2*^ = 20.81, *d*.*f*. = 1, *p* < 0.001). Conidiospore inoculated ants with their MG-sealed in group conditions survived significantly longer from those in solitary conditions (**Fig 3**). Metapleural gland sealing did not make any difference on the ant survival, with sham inoculations regardless of the environment (*χ*^*2*^ = 2.19, *d*.*f*. = 1, *p* = 0.139).

Mixed group treatments with sham and pathogen exposed ants resulted in following: a group of 5 spore inoculated and 8 sham inoculated had a lower survival than ants in a group of all sham inoculated (χ2 = 21.57, d.f. = 1, p <0.001) and a higher survival than the ants in a group of all spore inoculated ants (χ2 = 17.63, d.f. = 1, p <0.001). Under the assumption that only spore inoculated ants died within the duration of experiment, the spore inoculated ants in mixed inoculation group conditions had a surviving proportion of 0.47±0.13 (*prop±CI*), which was not different from ants in all inoculated group conditions that had a surviving proportion of 0.44±0.07 (*prop±CI*) (*two-tailed, z*=0.288, *p* = 0.772, suggesting that functional MG is necessary for individual survival but not so much for the nestmate survival (**Fig 3**).

## Discussion

To offer insight into social immunity and its importance for social insect species, we have tested how the group living or ant isolation, with or without functional MG, influences ant’s survival to pathogen exposure. We observed that nestmate presence was of high importance for the survival of ants exposed to fungal pathogen, where inoculated ants in isolation suffered much lower survival than those in groups. The MG appears to be crucial for the pathogen resistance of individual ants where those with sealed glands and inoculated with the fungus had little chance of survival over twelve days. However, a sealed MG did not appear to alter social survival effects; with a sealed MG, those in groups still had a higher survival probability than those in isolation. Overall, we present evidence that, with *T. curvispinosus*, nestmate presence improved disease resistance, implicating that the benefits of social immunity outweigh the increased risk of pathogen spread. Along with this, we find that the MG plays a crucial role in antisepsis and overall survival from pathogen inoculation.

Demonstrating an increase in pathogen resistance when living in groups has major implications to the collective idea of social immunity. While social immunity encompasses many preventative behaviors such as avoidance and exclusion [16], we show that social behaviors, such as grooming, are also important for increasing pathogen resistance for a nestmate rather than only excluding those infected. Grooming to remove infectious particles has been shown to improve pathogen resistance in other social species, such as in termites [39] and other ant species [40]. Isolation decreasing survival probability due to factors beyond the pathogen’s effect is unlikely over the time span of the experiment. There was a high survival in all sham inoculated treatments and no difference between group and isolated ants when lacking *M. brunneum* inoculation. Our results are consistent with earlier findings on *Acromyrmex* leaf-cutting ants, in which survival from fungus *M. anisopliae* inoculation was decreased when in isolation compared to grouped living [41]. We also note that fungal inoculated ants did not seem to spread to non-inoculated nestmates within the same group. The survival probability of the group containing only 5 of 13 inoculated ants had a survival probability consistent with a mortality rate of about 3.25 ants.

In terms of individual pathogen resistance, the MG appears to serve a crucial role in the ant’s survival probability. These results support the antisepsis hypothesis, suggesting that the MG secretes antimicrobial compounds to suppress pathogens [31]. The decrease in survival probability for infected *T. curvispinosus* with a sealed MG gland is consistent with what has been observed in leafcutter species, though not all species tested showed similar results [34]. Along with individual pathogen resistance, we suspected that the MG has an important part in reducing pathogen susceptibility in nestmates, contributing to the social immunity of the whole colony. Indeed, previous studies have shown that a functional MG gland has implications on overall colony fitness by improving brood survival rate when challenged with a pathogen and sanitizing nest materials [42]. In our study, both a group environment and a functional MG appear to be important for the pathogen resistance of *T. curvispinosus*, we did not find that a loss of MG function counteracted the improved survival probability from a group environment.

Larger colonies may have more reliance on the MG [34], whereas *T. curvispinosus* colonies often containing under 100 workers. Previous research has also suggested more general importance of MG for fungal growing ant species, due to stronger pathogen pressure [43]. Perhaps removing MG gland function within a larger, fungal-growing species, would show stronger effects to social pathogen resistance. With *T. curvispinosus*, infected individuals may rely on combining MG secretions for themselves and allogrooming from nestmates depending on the pathogen in question [44]. Species that have evolved a loss of MG, notably weaver ants, further complicate the balance between MG reliance compared to other behaviors such as grooming. Contrary to expectations, weaver ants did not have a lower survival probability than species with functional MG glands when exposed to a pathogen, and in addition appear to have an increase in grooming behavior [36].

Overall, our data suggest that group living is central for pathogen resistance in *T. curvispinosus* ants, supporting and expanding on the collective idea of social immunity within social insects. Mutually, the MG also serves a vital function in individuals’ survival when exposed to *M. brunneum*. While reducing survival probability, loss of MG function does not necessarily remove the survival benefits of grouped living, possibly indicating more reliance on behaviors such as allogrooming. Genomic and RNA analysis could offer more insight into the differences in infected ants in isolation compared to within a group. Behavioral and network analysis could also help model the reliance on behaviors compared to MG secretions among species. In general, exploration of social immunity can expand into the broad evolution of immunity and the evolutionary pressures important for transitioning from solitary to social structures [15]. It has recently been suggested that the glandular secretions of ants might have antibiotic and antifungal potential to substitute conventional drugs [46]. The identification of specific active chemical compounds secreted by *T. curvispinosus* metapleural glands that improve individual survival may be of significance in developing such methods.

## Acknowledgments

We thank T.A. Linksvayer for the input during experimental design. We thank Rowan University, College of Science and Mathematics SEED funding awarded to Vojvodic Kruse.

## Notes

### Competing Interest Statement

The authors have declared no competing interest.

